# Top-down contribution to motor network reorganization during action preparation

**DOI:** 10.1101/2021.12.20.473439

**Authors:** Alberto Pisoni, Valentina Bianco, Eleonora Arrigoni, Francesco Di Russo, Leonor J. Romero Lauro

## Abstract

**BACKGROUND:** It is unclear whether the Bereitschaftspotential (BP) recorded in humans during action preparation mirrors motor areas activation escalation, or if its early and late phases reflect the engagement of different functional networks.

**OBJECTIVE:** Here, we aimed at recording the TMS evoked-potentials (TEP) stimulating the supplementary motor area (SMA) to assess whether and how cortical excitability and functional connectivity of this region change as the BP increases. We hypothesize that, at later stages, the SMA functional network should become more connected to regions relevant for the implementation of the final motor plan.

**METHODS:** We performed TMS-EEG recordings on fourteen healthy subjects during the performance of a visuomotor Go/No-go task, eliciting and recording cortical activity and functional connectivity at -700 ms and -300 ms before the onset of visual stimuli over the SMA.

**RESULTS:** When approaching stimulus onset, and thus BP peak, the SMA increased its functional connectivity with movement-related structures in the gamma and alpha bands, indicating a regional top-down preparation to implement the motor act. Beta-band connectivity, instead, was maintained constant for the whole BP time-course, being potentially related to sustained attention required by the experimental task.

**CONCLUSION:** These findings reveal that the BP is not a mere result of increased activation of the SMA, but the functional networks in which this region is involved qualitatively changes over time, becoming more related to the execution of the motor act.

## Introduction

The Bereitschaftspotential (BP) is a slow negative cortical potential preceding the onset of voluntary movement, reflecting an increase in cortical excitability of the central region during motor preparation [1]. Considering self-paced and externally triggered [2,3] movements of the upper limbs, BP has been traditionally divided into two phases: the earliest phase (i.e., early BP or BP) starts approximately 1-2 seconds before voluntary movement and slowly rises over frontocentral regions with a symmetrical distribution; during the later phase, the potential suddenly increases its gradient around 500 ms before movement onset, displaying a steeper negative slope (i.e. late BP) and reaching a maximal amplitude (i.e. motor potential) over the contralateral central area [1]. However, it is unknown if this event relates to the increased activity of the motor network, or if a functional reorganization occurs as action execution, and thus the late phase, approaches. The recruitment of associative areas and other brain regions involved in higher-level cognitive processing becomes even more relevant in the wider context of action preparation [4], with distributed networks of cortical and subcortical regions playing a crucial role in regulating its different stages, including decision-making processes or the prediction of forthcoming events taking place in the surrounding environment [5–10].

Converging evidence coming from neuroimaging and brain stimulation studies [11–17] provided useful information on the main generators of the BP: the initial segment of the early BP is assumed to be generated by the bilateral activation of the supplementary motor area (SMA) and the cingulate motor area (CMA), thereafter the progressive increase in the potential’s amplitude likely reflects the additional involvement of the lateral premotor and, in the late phase, of primary motor areas, thus possibly reflecting the increasing interactions between the fronto-medial structures and the sensorimotor regions [18–20].

However, two main aspects of these preparatory motor ERPs are still unexplored: the functional dynamics involved in their occurrence, and if there is a change not only in the quantitative aspects of cortical activity but also a qualitative variation of their underlying functional networks.

The present study aimed at exploring the changes in cortical dynamics underlying the BP spatiotemporal evolution using an integrated transcranial magnetic stimulation (TMS) and electroencephalography (EEG) system.

TMS-EEG co-registration allows to probe real-time cortical reactivity through the analysis of TMS-evoked potentials (TEPs). TEPs are considered a reliable measure of cortical excitability, thus indexing the state of activation of the stimulated area [21,22]. Furthermore, TEPs analysis allows to observe the spatiotemporal propagation of the activity induced by TMS spreading in remote regions throughout functionally relevant connections, providing information about cortical connectivity [23].

In this study, healthy participants performed a visual Go/No-go task across four TMS-EEG recordings in which left SMA and a control region (left extrastriate area) were stimulated. TMS pulse occurred at two different times before stimulus presentation, in correspondence to the beginning (i.e., -700 ms) and the peak (i.e., -300 ms) of the BP, while TEPs were recorded from 60 scalp electrodes. This set-up allowed to explore the differences in cortical excitability of the SMA during the BP time-course by comparing TEPs at the two Stimulus Onset Asynchronies (SOAs). At the same time, functional connectivity analysis performed at the source level complemented this knowledge by assessing the networks in which the SMA was involved during action preparation, exploring the functional meaning of the neurophysiological activity recorded before action onset.

## Materials and Methods

### Participants

Fourteen healthy, right-handed volunteers (8 females, mean (±standard deviation, SD) age = 24.3 ±2.2 years) took part in this study. They all had a normal or corrected-to-normal vision and no history of neurological, psychiatric, or other relevant medical condition. Each participant completed a safety screening questionnaire to exclude the presence of contraindication to TMS following the current TMS safety guidelines [24] and gave informed written consent prior to their participation in the study. The study was performed in the TMS-EEG laboratory of the University of Milano-Bicocca in accordance with the Declaration of Helsinki and the approval of the local Ethics Committee.

### Procedures

Participants sat comfortably in a semi-reclined armchair in front of a 20” computer screen at a distance of 114 cm, with their arms on the armrests and their right hand positioned on a PC mouse, allowing them to press a button with their right index finger.

The experimental conditions were tested in a four-block recording session. In each block, participants were asked to perform an uncued, visual Go/No-go task during the TMS-EEG recording. Each trial started with a fixation point, a yellow dot displayed at the centre of the screen on a black background, which remained present on the screen for the entire trial. Then, in a random jittering interval ranging from 2900 ms to 3900 ms according to trial type, the TMS pulse was delivered, and after the corresponding SOA (i.e., 300 ms or 700ms) one of four different visual stimuli, made by four squared configurations with vertical and horizontal bars (4°x4° of visual angle, Fig.1), was randomly presented in focal vision for 250 ms, with equal probability (p=0.25). Two configurations were defined as targets, thus requiring the participants to produce the motor response (i.e., *Go* trials, p=0.5), and two as non-targets and they were associated with the instruction to withhold the response (i.e., *No-go* trials, p=0.5). Then, participants’ responses were collected (max 1500ms). In total, 60 trials for each experimental condition were collected (*Go*/*No-go* trials * 2 SOAs and * 2 cortical targets) for a total of 480 trials divided in 120 trials blocks. Additional 120 trials were recorded without TMS in another block. Blocks duration was approximately 10 minutes, and their order was counterbalanced between subjects.

**Figure 1.**
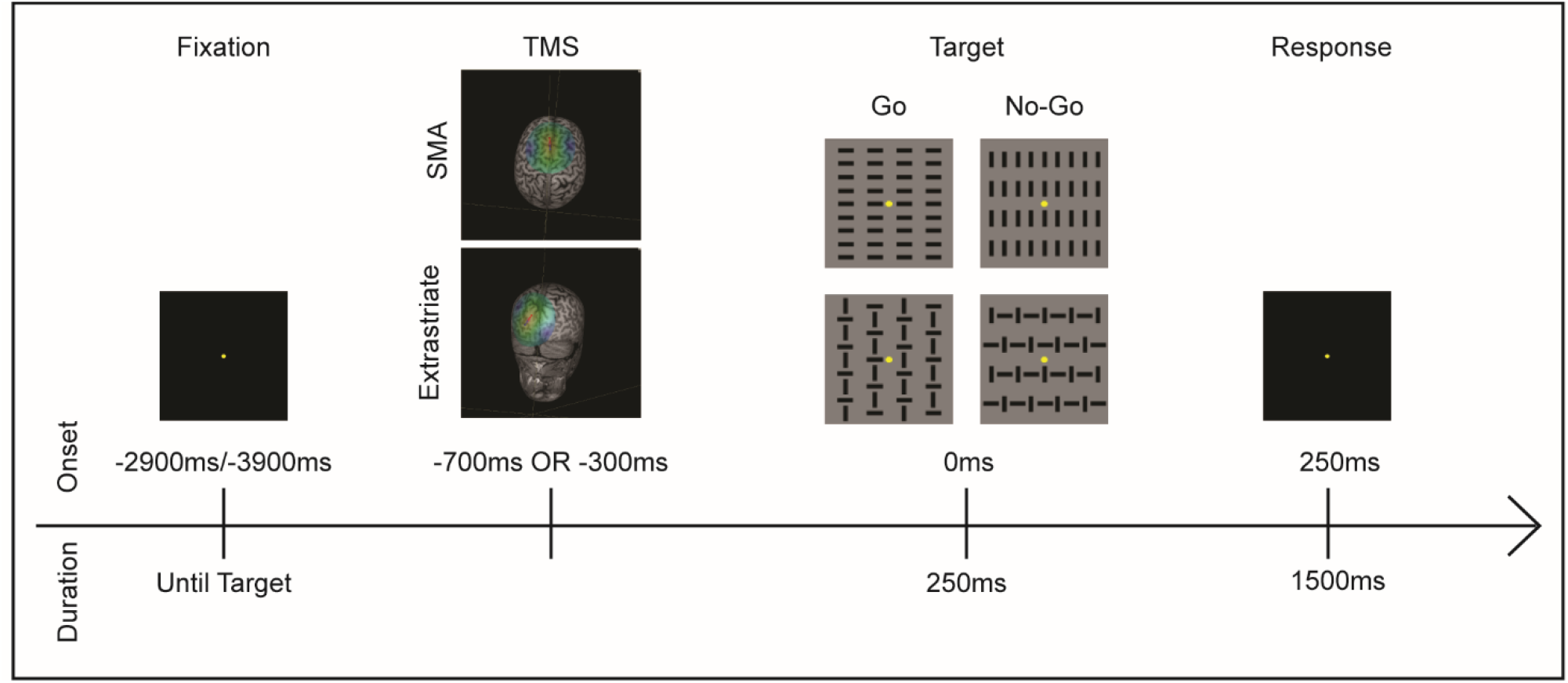
Timeline of an experimental trial. Trials started with a fixation screen, presented for a jittering interval between 2900 ms and 3900 ms. Within this interval, according to SOA condition, TMS was delivered -700 or -300 ms before Target onset on the left SMA or left extrastriate region, according to the experimental block. The Target appeared for 250 ms. Go and No-Go trials appeared with equal probability (p=0.05), with each figure appearing 25% of the trials. Finally, a fixation screen was displayed to collect participants’ response. In No TMS blocks, the experimental procedure remained the same but TMS was not delivered.

TMS pulses were delivered at two alternative scalp sites (2 blocks each), following MNI coordinates derived from a previous fMRI study [25]: the left SMA (−4, 17, 45) a potential generator of the BP, and the left extrastriate visual cortex (−21, -98, 5), unrelated to BP origins. Before the experimental session, a short practice familiarized participants with the stimuli and the task.

Trials randomization, TMS and visual stimuli timing, and behavioural data recording (i.e., response times, RT, and accuracy) were under computer control (E-Prime, Psychology Software Tools Inc.), as well as TMS and EEG trigger delivery.

### TMS stimulation

Single-pulse biphasic TMS was delivered with an Eximia™ TMS stimulator (Nexstim™, Helsinki, Finland) using a biphasic focal figure-of-eight 70mm coil. The stimulation target sites were localized on normalized individual high-resolution (1 mm^3^) MRI images at the selected MNI coordinates. The location of the two stimulation targets was identified for each participant using a Navigated Brain Stimulation (NBS) system (Nexstim™, Helsinki, Finland) based on infrared-based frameless stereotaxy, allowing also an accurate monitoring of the position and orientation of the coil and an online estimation of the distribution and intensity (V/m) of the intracranial electric field induced by TMS. TMS intensity was preliminarily adjusted for each participant and cortical target before the experiment, to ensure a cortical response of at least 6 µV, as assessed online in a short recording session before the experimental blocks. The mean intensity of the electric field induced by TMS was 98.57 V/m (SD=±16.55) for the left SMA, and 94.86 V/m (SD=±17.18) for the left extrastriate region, corresponding to a mean intensity – expressed as percentage of the maximal stimulator output (MSO) – of 63.21% (SD=±2.49) for the SMA, and 63.21% (SD=±2.49) for the extrastriate region.

### EEG recording and pre-processing

EEG data were continuously acquired from 60 channels using a sample-and-hold [26] TMS-compatible system (Nexstim™, Helsinki, Finland). Two electrodes were placed over the forehead as ground. Two additional electro-oculographic (EOG) channels were placed near the eyes and used to monitor ocular artifacts due to eye movements and blinking. Noise-masking was performed by continuously playing into earplugs an audio track created by shuffling TMS discharge noise, to prevent the emergence of auditory evoked potentials (as in [27–29]). Electrodes’ impedance was kept below 5 kΩ. EEG signals were acquired with a sampling rate of 1450 Hz.

Data pre-processing was carried out using Matlab R2019a (Mathworks, Natick, MA, USA). First, EEG data was down-sampled to 725 Hz. Continuous signal was segmented in epochs starting 1000 ms pre- and ending 1000 ms post-TMS pulse. A band-pass filter between 0.2 and 80 Hz and a 50 Hz notch filter were applied to the selected epochs. Single trials with excessive artifacts were rejected by visual inspection. Bad channels were excluded and then interpolated using a spherical interpolation function included in EEGLAB [30].

TMS-evoked potentials (TEPs) were computed by averaging artifact-free trials, re-referenced using an average reference, and baseline-corrected between -900 and -700 ms before TMS pulse. Independent Component Analysis (ICA) was performed to remove residual artifacts due to TMS pulse, muscle contraction, or eye movements.

### Experimental design and data analysis

As the aim of the study was to investigate the cortical dynamics underlying action preparation, all the analyses focused exclusively on the pre-stimulus time window. Therefore, the distinction between *Go* and *No-Go* Trials was not taken into account [10].

The experimental design was a factorial 2×2 design, with SOA (2 levels: -700 and -300ms) and target area (2 levels: left SMA and left extrastriate region).

First, to confirm the BP occurrence in the present paradigm, for the No-TMS block the ERPs associated with the visual stimulus onset were calculated. For this purpose, EEG was segmented in 1500 ms windows beginning 1200 ms before visual stimulus onset and ending 300 ms after.

Then, data from TMS blocks was analysed using the Fieldtrip toolbox [31]. First, to track the changes in cortical excitability occurring during the BP time-course, we compared TEPs amplitude at the two TMS SOAs (i.e., -700 ms vs -300 ms) within each stimulation site (i.e., SMA and extrastriate). The comparison between the two SOAs was restricted within two electrode clusters of interest selected in correspondence to the stimulation sites: SMA cluster included F1, FC1, FCz, C1, Cz; extrastriate cluster included P1, Pz, PO3, POz, O1, Oz. This choice was supported by previous literature on the BP, as the component is mainly detectable at centro-medial sites (e.g. Cz, as in [8]). Within each cluster, TEPs amplitude was compared using a dependent sample t-test with a nonparametric cluster-based permutation approach. This procedure tests the differences between two experimental conditions by taking into consideration the spatiotemporal structure of EEG data, thus allowing to correct for multiple comparisons by permuting the data and clustering them based on their spatial and temporal proximity [32]. In the present study, for each comparison, we performed 10000 permutations on the time window between 0 and 250 ms post-TMS and used a permutation-significance threshold of p=0.05. Using the same approach, two additional analyses were carried out comparing between the two SOAs the frontal evoked response (i.e., within SMA cluster) during extrastriate stimulation and the occipital evoked response (i.e., within the extrastriate cluster) during SMA stimulation.

For SMA trials, functional connectivity was computed at the source level. The forward model was created starting from a Boundary Element Model (BEM) obtained segmenting a subject MRI into five standard tissues (Gray and white matters, CSF, Skull, and Scalp). The head model was then computed assigning standard conductivity values for the scalp, skull, and brain compartments [33,34]. Source space was defined performing a cortical reconstruction and volumetric segmentation of the grey matter with the Freesurfer image analysis suite [35], down-sampled to 8193 cortical sources, and realigned to the head model space. Finally, individual lead-field matrices were computed by aligning this forward model with the individual electrode positions, which were recorded during each TMS-EEG session. This model was also used to compute the spatial filter matrix, i.e., the inverse of the lead field matrix. Source reconstruction of the EEG time-series was conducted with the eLORETA method implemented in Fieldtrip [36–38] for solving the inverse modelling. Source signals were then segmented in 88 regions of interest (ROIs) according to the AAL atlas [39]. The computed source time-series were then averaged to index each ROI source activation [40] and functional connectivity was performed. Specifically, Phase Locking Value (PLV, [41,42]) was computed between the left SMA and the other brain parcels for alpha (8-12 Hz), beta (13-30 Hz), gamma (31-40 Hz) bands for the two TMS SOAs, with a particular interest for beta and gamma frequency, as suggested by Kim et al. ([43]). This analysis was also performed on a surrogate dataset created by shuffling the phase of the source reconstructed time series for each experimental condition. To reduce the risk that spurious connectivity could be included in our results, real data were compared with the surrogate ones computing a t-test performed on each connectivity pair (87, SMA with the other 87 brain parcels) and corrected for multiple comparisons based on 2000 permutation approach, implemented in Matlab, with a significance level of p=0.001 [29]. Surviving connections were plotted to highlight the resulting FC between the left SMA and the other parcels. Finally, to compare the resulting connectivity patterns for each frequency band across experimental conditions, the network strength, computed as the sum of the PLV of the resulting connections, was pairwise compared against a null distribution created using a permutation approach [44].

## Results

### Behavioural results

Participants had a mean accuracy in detecting the go signals of 98.1% (SD=±2.3), 98.1% (SD=±2.6), and 97.2% (SD =±3.4) in the No TMS, SMA, and extrastriate stimulation blocks respectively. False alarms were 3.6% (SD=±5.1) 4.8% (SD=±3.4) and 5.7% (SD=±6.5) respectively, and RTs were 588 ms (SD=±48), 581 ms (SD=±44), and 579 ms (SD=±33). These values are in line with the literature using a comparable inter-stimulus interval [25], and, as expected, showed no difference across No-TMS, SMA, and extrastriate stimulation recordings (t-tests, all ps>0.13). The high accuracy in performing the Go/No-go task ensures the attention of the participants to the experimental procedures.

### Pre-stimulus ERPs

The ERP waveforms and scalp topography related to the visual stimulus onset obtained in the No-TMS condition are shown in Fig. 2. The BP was detectable with an onset at -950 ms as a gentle increase just before the stimulus onset, reaching a maximal amplitude of 4 μV. In this period, the BP had a consistent scalp topography focusing on medial centro-parietal areas. This result confirms the presence of BP in the present paradigm.

**Figure 2.**
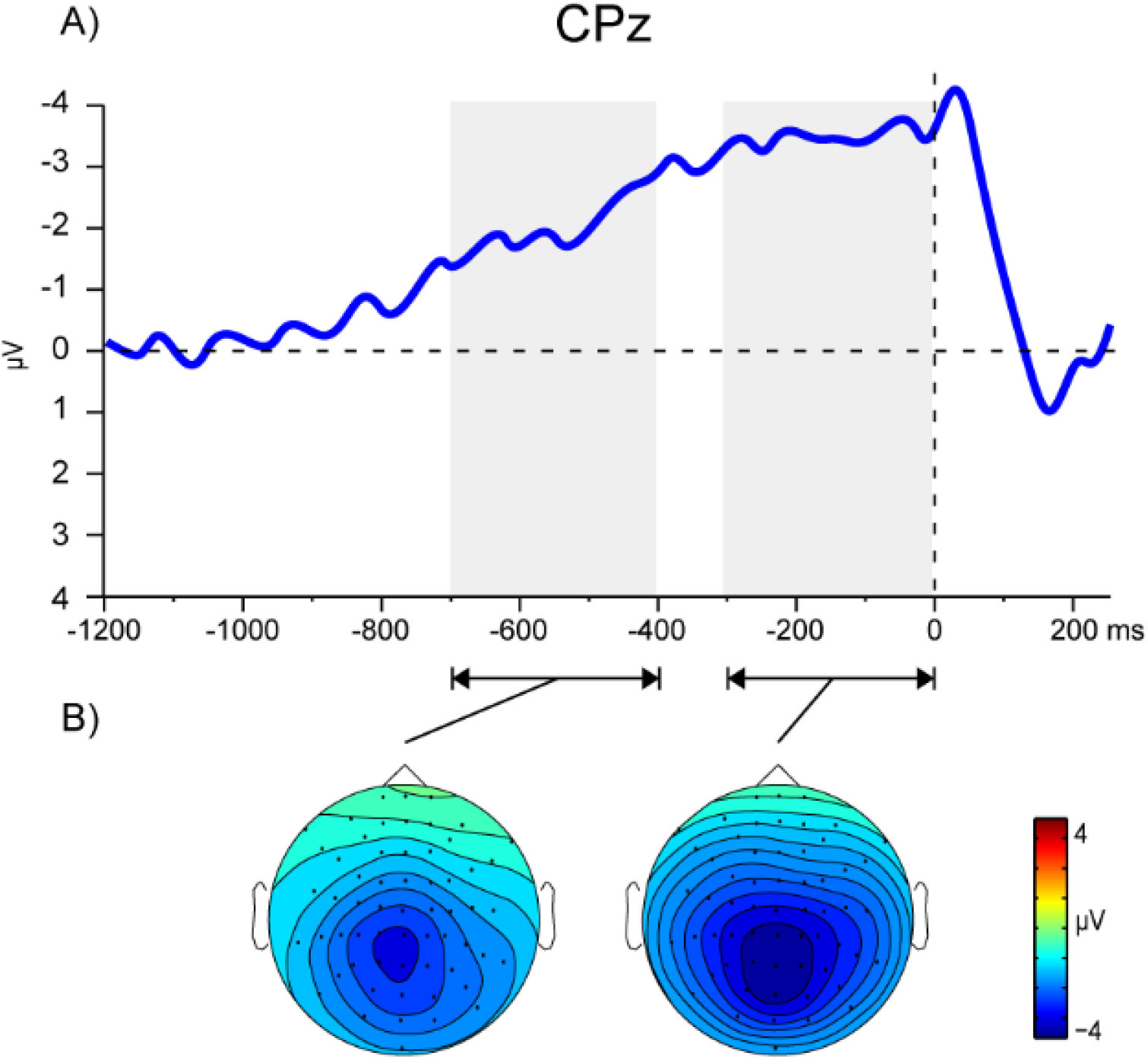
Pre-stimulus ERP in the No-TMS condition. The BP time-course (A) and its scalp topography in two windows (B, from -700 to -400 and from -300 to 0 ms) representing the early and late BP phases are displayed.

### Cortical reactivity during the BP time-course

To assess the changes in cortical excitability during BP time-course, we compared the amplitude of TMS-evoked activity between SOAs (i.e., -700 ms and -300 ms) within each stimulation condition (i.e., left SMA and extrastriate) employing a cluster-based permutation t-test on the 0-250 ms post-TMS time window [32].

Fig. 3a shows the butterfly plot of the grand average of the TEPs evoked by SMA stimulation at the -700 ms (upper row) and -300 ms (lower row) SOAs. For both the SOAs, the TMS pulse produced seven main components, peaking respectively around 5, 10, 20, 45, 65, 100, and 150 ms. The first three components do not show clear differences between the two SOAs. Conversely, the component peaking at 45 ms presents a greater amplitude when TMS was applied at the -300ms SOA. Statistical comparison of the two SOAs confirmed these observations. A cluster-based t-test revealed a difference between the two SOAs in correspondence of a negative cluster between 35 and 50ms post-TMS (p=0.0058), corresponding to the negative component peaking at 45 ms (Fig. 3c). Scalp topographies of the N45 component (Fig. 3b) showed in both SOAs conditions a negative focus located at medial centro-frontal sites more prominent at the left hemisphere, corresponding to the stimulated area.

**Figure 3.**
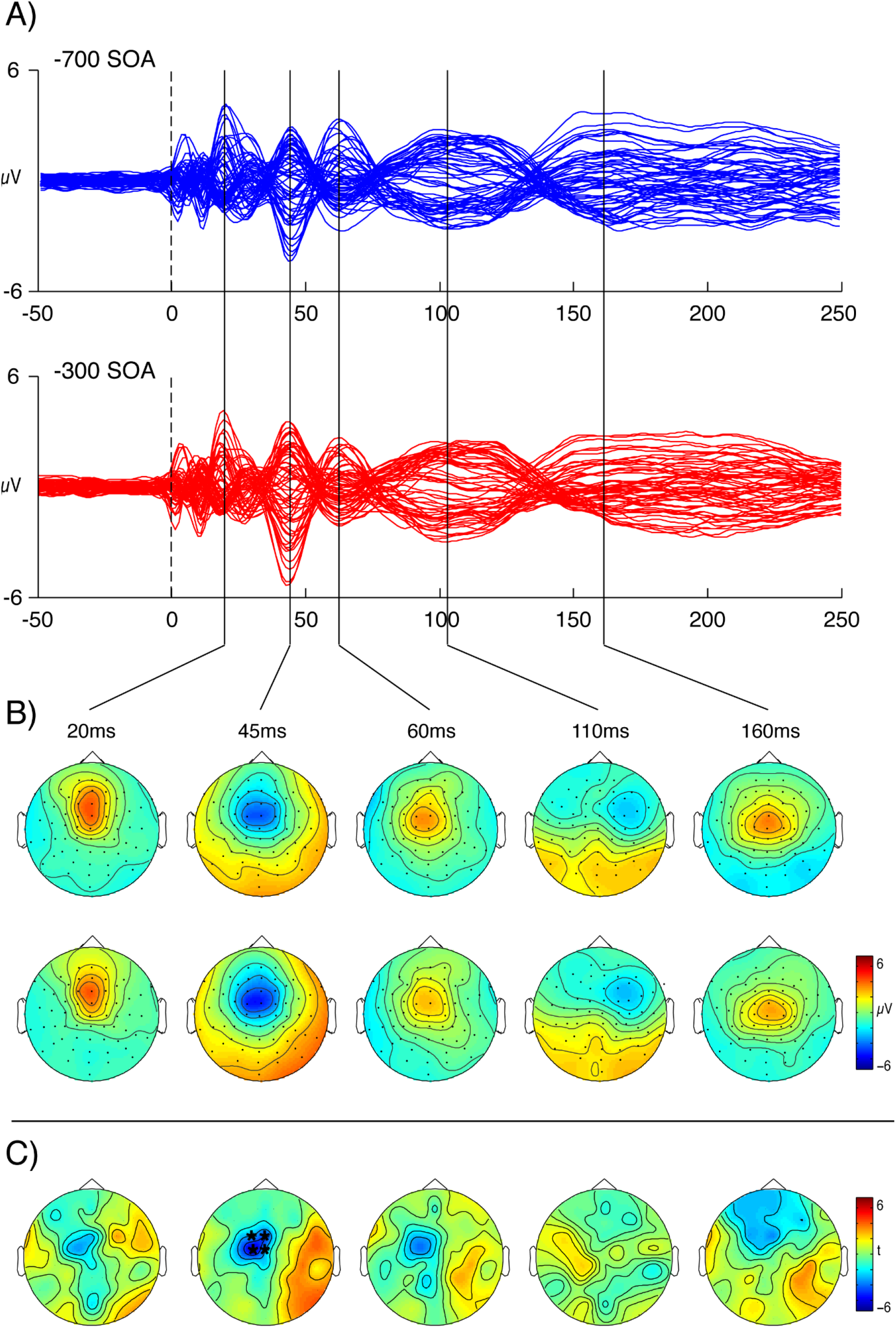
Grand average results from TEPs analyses. (A) Upper row, butterfly plots of TEPs recorded in -700 ms SOA trials (blue lines). In the lower row, butterfly plots of TEPs in the -300 ms SOA condition are reported (red lines). (B) Scalp topographies of five components peaking around 20 ms, 45 ms, 60 ms, 110 ms and 160 ms are reported for the -700 ms (upper row) and -300 ms (lower row) SOAs. (C) Scalp topographies of the five main TEP components representing t statistics between the -700 ms and the -300 ms SOA conditions. Channels highlighted by asterisks are significantly different in the two conditions as revealed by a cluster-based permutation analysis.

### Insert Figure 3 about here

Activity in the SMA ROI confirmed these observations (Fig. 4, upper left panel). Concerning the extrastriate region stimulation, the TMS-evoked activity showed no differences between the two TMS SOAs in terms of amplitude, latency, and spatial distribution (Fig. 4, lower right panel). Five components were observed, reaching their peaks respectively around 5, 10, 30, 80, and 150 ms. All the components were spatially distributed over the stimulated area (left parieto-occipital region), with no appreciable difference between SOAs. In line with these observations, no statistical difference was found between the two conditions.

**Figure 4.**
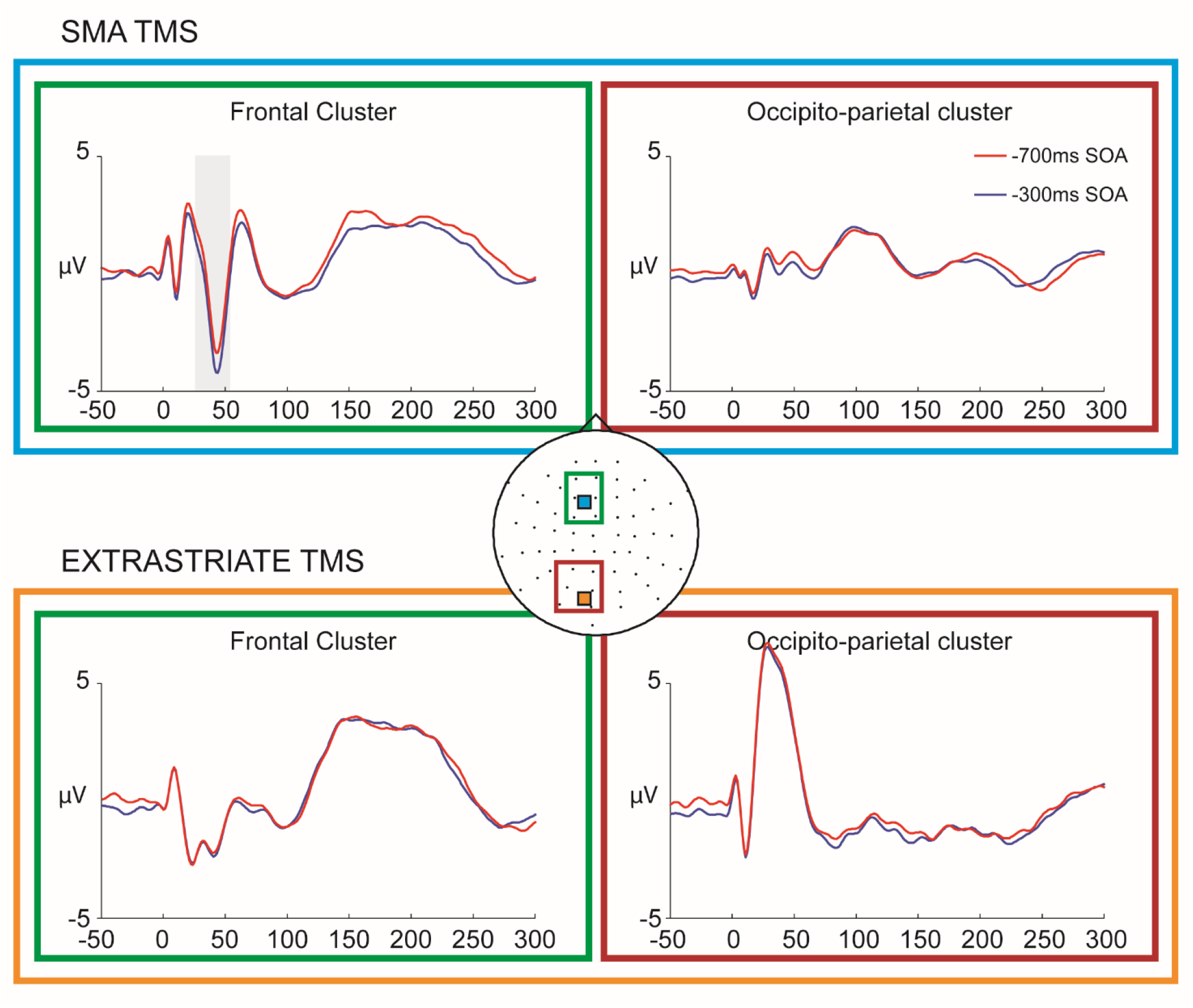
Grand average results from clusters TEPs analyses. Blue box: TEPs obtained in the frontal (green box, left) and parieto-occipital (red box, right) electrode clusters when stimulating the left SMA. Gray shaded areas represent significant differences between the -700 ms SOA (blue line) and the – 300 ms SOa (red line). Orange box: TEPs obtained in the frontal (green box, left) and parieto-occipital (red box, right) electrode clusters when TMS targeted the left extrastriate region

To control for the spatial specificity of our findings, we performed two additional analyses comparing at the two SOAs the left occipital evoked response during SMA stimulation (Fig. 4, upper left panel) and the left frontal evoked response during extrastriate stimulation (Fig. 4, lower left panel). In both cases, we found no significant differences.

### Functional networks during action preparation

Concerning functional connectivity analysis, we computed source-level the Phase-Locking Value (PLV, [41,42]) between the left SMA and the other cortical parcels within three frequency bands (i.e. alpha, beta and gamma), to make a comparison between SOAs.

Concerning the gamma band (31-40 Hz), TMS delivered over the left SMA at -700 ms before visual stimulus onset highlighted bilateral connections between the left SMA and fronto-medial and parietal structures, such as the right SMA, the left superior, and middle frontal gyri, the bilateral middle cingulate cortex and paracentral lobules, the right precuneus and postcentral gyrus (Fig. 5a, Tab 1). At -300ms SOA, the SMA increased its connectivity with frontal, parietal, and occipital regions, including bilateral connections to the precentral gyri, the superior frontal gyri, the middle cingulate cortex, the paracentral lobules, and the bilateral superior occipital gyri, the left inferior frontal gyrus and the superior parietal cortex, the right medial superior frontal gyrus (Fig. 5a, Tab. 1a). The increase in the total network strength was significant between the -700 ms and -300 ms SOAs (Fig. 5b).

**Figure 5.**
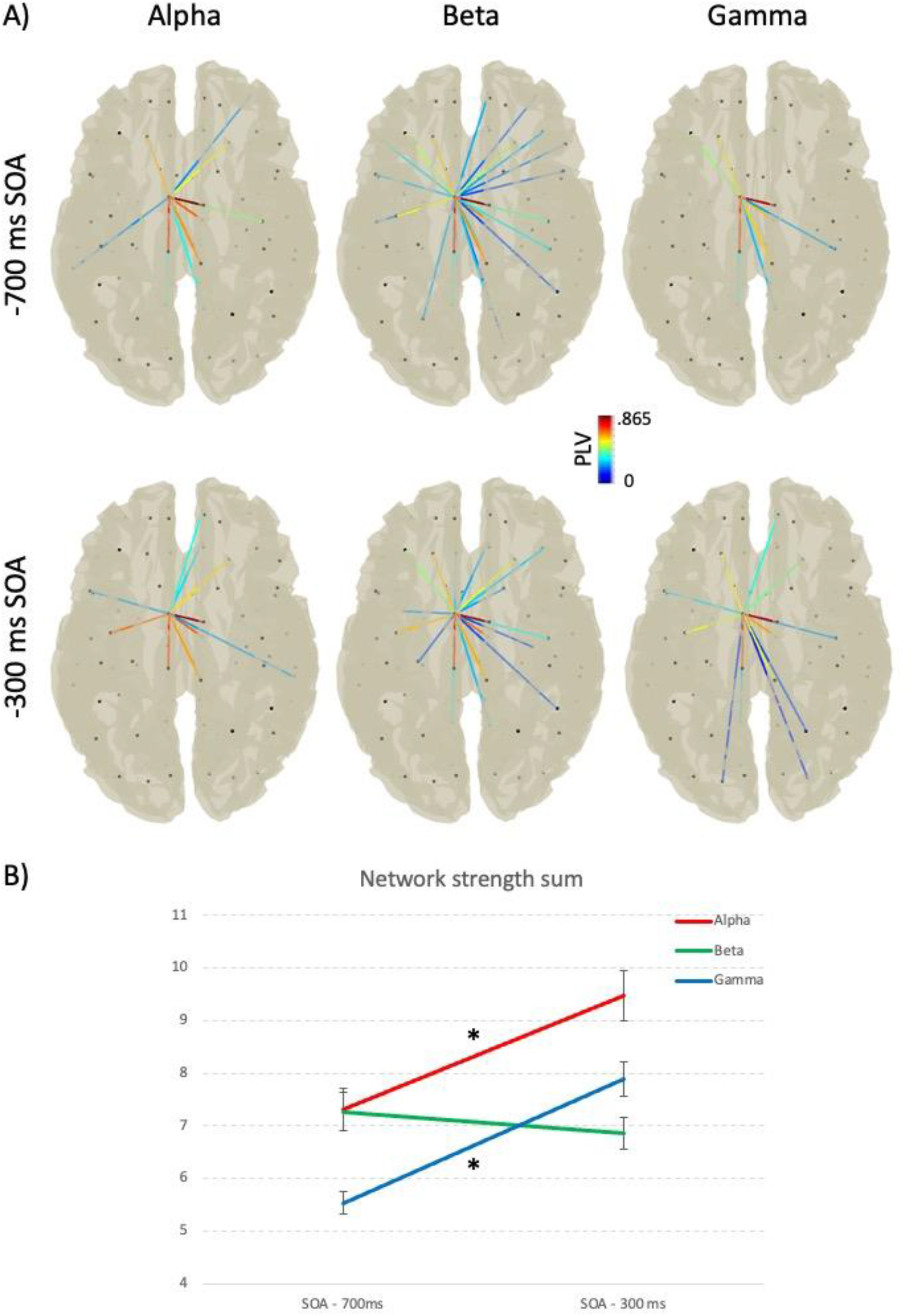
Source connectivity results. (A) Plots for the alpha (8–12 Hz) beta (13-30 Hz) and gamma (31-40 Hz) bands during SMA TMS sessions for the -700ms and - 300ms SOAs conditions are displayed. Black dots represent the centroids of the 88 parcellated cortical regions. Connections start from the left SMA area. Labels of the connected regions are reported in Tables 1 2 and 3. The color of the connection represent PLV values. (B) plot of the total network strength in the three different frequency bands and in the two SOA conditions. Asterisks represent significant differences between the two SOAs as resulting from the comparisons against a null distribution created using a permutation approach. Error bars represent +- 1 MSE.

Conversely, connectivity in the beta band (13-30 Hz) remained stable throughout the BP rising phase: comparing the networks between SOAs, we found that left SMA maintained its connections with several structures of both hemispheres, including the right SMA, the paracentral lobules, the cingulate cortex, and the precuneus bilaterally (Fig. 5a, Tab. 1b). The network strength in the beta band did not significantly change between the two SOAs (Fig. 5b).

**Table 1.** Results of the connectivity analysis. List of significant connections in the -700 ms and -300 ms SOAs in the Gamma (a), Beta (b) and Alpha (c) connectivity bands.

| **a) Left SMA gamma-band connectivity** |  |
| --- | --- |
| **-700 ms** | **-300 ms** |
| Left Superior Frontal Gyrus | Left Precentral |
| Left Middle Frontal Gyrus | Right Precentral |
| Right Supplementary Motor Area | Left Superior Frontal Gyrus |
| Left Middle Cingulum | Right Superior Frontal Gyrus |
| Right Middle Cingulum | Left Inferior Frontal Gyrus (pars opercularis) |
| Left Precuneus | Right Supplementary Motor Area |
| Left Paracentral Lobule | Right Superior Medial Frontal Gyrus |
| Right Paracentral Lobule | Left Middle Cingulum |
| Right Postcentral Gyrus | Right Middle Cingulum |
| Right Precuneus | Left Superior Occipital Gyrus |
|  | Right Superior Occipital Gyrus |
|  | Right Superior Parietal |
|  | Left Precuneus |
|  | Left Paracentral Lobule |
|  | Right Paracentral Lobule |
| **b) Left SMA beta-band connectivity** |  |
| **-700 ms** | **-300 ms** |
| Right Precentral Gyrus | Left Precentral Gyrus |
| Left Superior Frontal Gyrus | Right Superior Frontal Gyrus |
| Right Superior Frontal Gyrus | Right Supplementary Motor Area |
| Left Middle Frontal Gyrus | Left Rectus |
| Left Inferior Frontal Gyrus (pars opercularis) | Left Insula |
| Right Supplementary Motor Area | Left Middle Cingulum |
| Left Middle Cingulum | Right Middle Cingulum |
| Right Middle Cingulum | Right Lingual Gyrus |
| Right Postcentral Gyrus | Right Inferior Parietal |
| Left Precuneus | Left Precuneus |
| Right Precuneus | Right Precuneus |
| Left Paracentral Lobule | Left Paracentral Lobule |
| Right Paracentral Lobule | Right Paracentral Lobule |
| Right Putamen | Left Caudate |
|  | Right Caudate |
| **c) Left SMA alpha-band connectivity** |  |
| **-700 ms** | **-300 ms** |
| Left Precentral Gyrus | Left Precentral Gyrus |
| Right Precentral Gyrus | Right Precentral Gyrus |
| Left Superior Frontal Gyrus | Right Superior Frontal Gyrus |
| Right Superior Frontal Gyrus | Right Superior Orbital Frontal Gyrus |
| Left Middle Frontal Gyrus | Left Inferior Frontal Gyrus (pars opercularis) |
| Right Middle Orbital Frontal Gyrus | Right Supplementary Motor Area |
| Right Supplementary Motor Area | Left Olfactory |
| Left Middle Cingulum | Left Medial Orbital Frontal Gyrus |
| Right Middle Cingulum | Right Medial Orbital Frontal Gyrus |
| Left Supramarginal Gyrus | Right Rectus |
| Left Precuneus | Left Insula |
| Right Precuneus | Right Anterior Cingulum |
| Left Paracentral Lobule | Left Middle Cingulum |
| Right Paracentral Lobule | Right Middle Cingulum |
|  | Left Postcentral Gyrus |
|  | Right Supramarginal Gyrus |
|  | Left Precuneus |
|  | Left Paracentral Lobule |
|  | Right Paracentral Lobule |
|  | Right Putamen |
|  | Left Pallidum |

Connectivity in the alpha range (8-12 Hz) significantly increased with the BP time-course. At -300 ms SOA (Fig. 5a), new interactions arose between the left SMA and several regions in the frontal lobe, such as the right superior frontal cortex, the left inferior frontal cortex, the right anterior cingulate cortex, and the left insula. Moreover, the left SMA appears to be connected to posterior parietal regions and the basal ganglia (Tab.1c). This increase in total network strength was statistically significant (Fig. 5b).

## Discussion

Our findings shed light on the activity and functional organization of motor networks during action preparation. Critically, in line with previous literature, we confirmed the crucial role of the SMA in the generation of the BP: when TMS is delivered over the SMA in the later BP phase, greater cortical activity is elicited in the form of a steeper negative response. Since early TEP components, evoked in the first 50 ms post-TMS, mainly reflect a local response to stimulation, coming from the cortical sites near the targeted region [27,28], it follows that the SMA increased its activity concurrently to BP rising phase, and therefore, the reported difference in the response magnitude in frontal responses between the two SOAs may be considered as a direct proof of the increased cortical excitability of the stimulated region during later stages of preparation. Accordingly, we reported no significant differences by comparing SMA-related TEPs in a distant electrode cluster, namely in the parieto-occipital region, or stimulating a control area (the extrastriate region).

Source modelling applied to TMS-EEG data allowed us to highlight significant connections between the left SMA and other regions that are known to be crucially involved in motor planning and action preparation, with complex patterns of functional reorganization in cortical networks related to movement implementation. Our findings support the relevance of gamma, beta, and alpha distributed networks which might play a role in the processing of specific cognitive computations at early stages of movement preparation [43,45,46].

Critically, we report an increase in gamma-band connectivity, which was suggested to support a regional, short-range communication that is crucial for the local processing of multiple segregated inputs coming from different areas [46–48]. Gamma synchronization is indeed considered a key mechanism underlying precise, selective, and effective communication within activated networks, by orchestrating the activity of neuronal assemblies located in different brain regions but functionally related and engaged in the execution of a cognitive task [49,50]. Moreover, gamma synchronization has been related to a pro-kinetic role in the generation of voluntary movement (e.g., [51–53]). Here, an early, bilateral communication between the SMA and other cortical regions belonging to the medial frontal cortex, particularly involved in the cognitive aspects of action preparation and control was found, suggesting a top down potential modulatory action on motor functions [54–57]. Previous evidence, indeed, suggested that interactions between the middle cingulate cortex (MCC) and SMA might play a pivotal role in keeping the activity sustained during BP time-course, until the movement is finally triggered via the activity of the primary motor cortex [58].

In correspondence to the late BP, instead, the SMA was significantly connected with sensorimotor structures crucially involved in movement initiation and execution [59], indicating a clear role in action implementation of this late functional network. Finally, by the time stimulus presentation approaches, SMA-related gamma-band communication widens to several posterior regions relevant for perceptual and attentional processing, as the superior occipital gyri, suggesting an increase in the information flow between the frontal, executive brain areas, responsible for the selection and implementation of the motor output, and sensory brain areas, responsible for the processing and evaluation of the upcoming visual imperative stimulus.

Additionally, SMA-related alpha-band connectivity also increased as the BP reached its peak. Alpha-band synchronization has been related to visual attention [60] and top-down neural networks responsible for attentional features crucial for task demands [61,62]. Previous evidence highlighted that regions in the dorsomedial frontal cortex could directly modulate alpha-band activity in the posterior visual areas, modulating visual stimuli sensitivity [63], thus subserving top-down modulation of sensory activity [64].

Beta-band connectivity, instead, resulted strong and stable throughout the whole BP rising phase, as network strength did not significantly differ between -700 ms and -300 ms SOAs. The functional meaning of beta-band activity can be seen in light of sensorimotor and cognitive information maintenance within distributed networks [65]. Beta synchronization has been closely related to top-down processing [66,67] and might sub-serve the active maintenance of information within large-scale networks in the absence of external stimulation [65,68]. Thus, beta-band connectivity might be related to proactive attentional control, as participants were required to actively monitor the appearance and features of visual stimuli to decide whether to act or not.

As a cautioning note, results have to be seen in the light of potential ghost connections issues, which are related to EEG and MEG interareal connectivity measures [69]. Moreover, our experimental design did not allow to assess the link between the observed inter-areal interactions and the behavioural performance, to elucidate the functional role of observed networks during action preparation.

The present study proves TMS-EEG as an efficient tool to dissect and track functional connectivity changes occurring during action preparation. To the best of our knowledge, this work presents for the first time novel TMS-EEG connectivity data shedding light on BP generation time-course and propagation during preparation stages of action, and provides unequivocal evidence of involvement of complex interareal networks aiming at proactively ensuring proper unfolding of motor acts. By supporting integration of functionally-related information into unitary representations and their active maintenance within distributed networks, high-frequency interactions might play a role in implementation of specific cognitive computations, which are determinant for specification of the motor program over the course of pre-stimulus stage [43,45,70].

Increased inter-areal communication during preparatory stages might be beneficial for setting the system for readiness before sensory events are presented. Then, after stimulus perception and categorization, decision making processes can take place and determine the appropriate motor output. Considering the interplay between inter-areal functional coupling and the regional activation of the motor system [71], the increased phase synchronization between cortical and subcortical brain structures may exert a modulatory action on brain regions activations, facilitating selection of appropriate motor output after stimulus presentation.

**The authors declare that they have no affiliations with or involvement in any organization or entity with any financial interest, or non-financial interest in the subject matter or materials discussed in this manuscript**.

## Funding

This work was supported by a grant from the University of Milano-Bicocca to AP (grant number 2018-ATE-0314).

## Author contributions

Conceptualization: AP, VB, FDR, LJRL Methodology: AP, EA, VB, FDR, LJRL

## Notes

### Competing Interest Statement

The authors have declared no competing interest.

